## Supplementary Figures for "EZHIP constrains Polycomb Repressive Complex 2 activity in germ cells"

**Supplementary Figures, Table and legends:**

Includes:

Supplementary Figure 1
Supplementary Figure 2
Supplementary Figure 3
Supplementary Figure 4
Supplementary Figure 5
Supplementary Figure 6
Supplementary Figure 7

Supplementary Table 1
Supplementary Table 2

Supplemental References.

### Supplementary Figure 1

(A) Western blot on nuclear extracts and Flag-IP from WT or EZH1-Flag mouse testes probed with anti-EZH1, Representative result, n=2. (B) Volcano plot representation of EZH1 interactome from EZH1-Flag mice testes IP compared to WT. In red are all the core complex subunits, in green the cofactors and in blue EZHIP, n=3. (C) Anti-FLAG western blot analysis on nuclear extracts from Hela-S3 cells stably overexpressing *Mus Musculus* or *Homo Sapiens* EZHIP and corresponding control; HDAC1 is used as a loading control, representative result. (D) Volcano plot representation of hEZHIP interactome after Flag-IP on HeLa-S3 overexpressing a tagged version of the protein. Same color code as in (B), additional interactors in black, (n=3). (E) Phylogenetic tree representing EZHIP protein sequence across placental mammals (Phylogenetic Analysis by Maximum Likelihood (PAML) algorithm). (F) PRC2 components and cofactors expression in mouse MII oocytes and α6+c-kit- spermatogonia, data extracted from the RNA-seq presented in Fig 4C and 5F (mean, n=2). (G) PRC2 components and cofactors expression in mouse somatic and germ cells [1] (GSE89711). m: male, f: female, PGC: primordial germ cells, SOMA: Somatic cells (mean, n= 3). (H) Expression of *EZHIP*, *PIWIL2*, *EZH2*, *JARID2* in human PGCs and somatic cells at week 7 [2] (NCBI SRA: SRP057098; mean, n≥ 3).

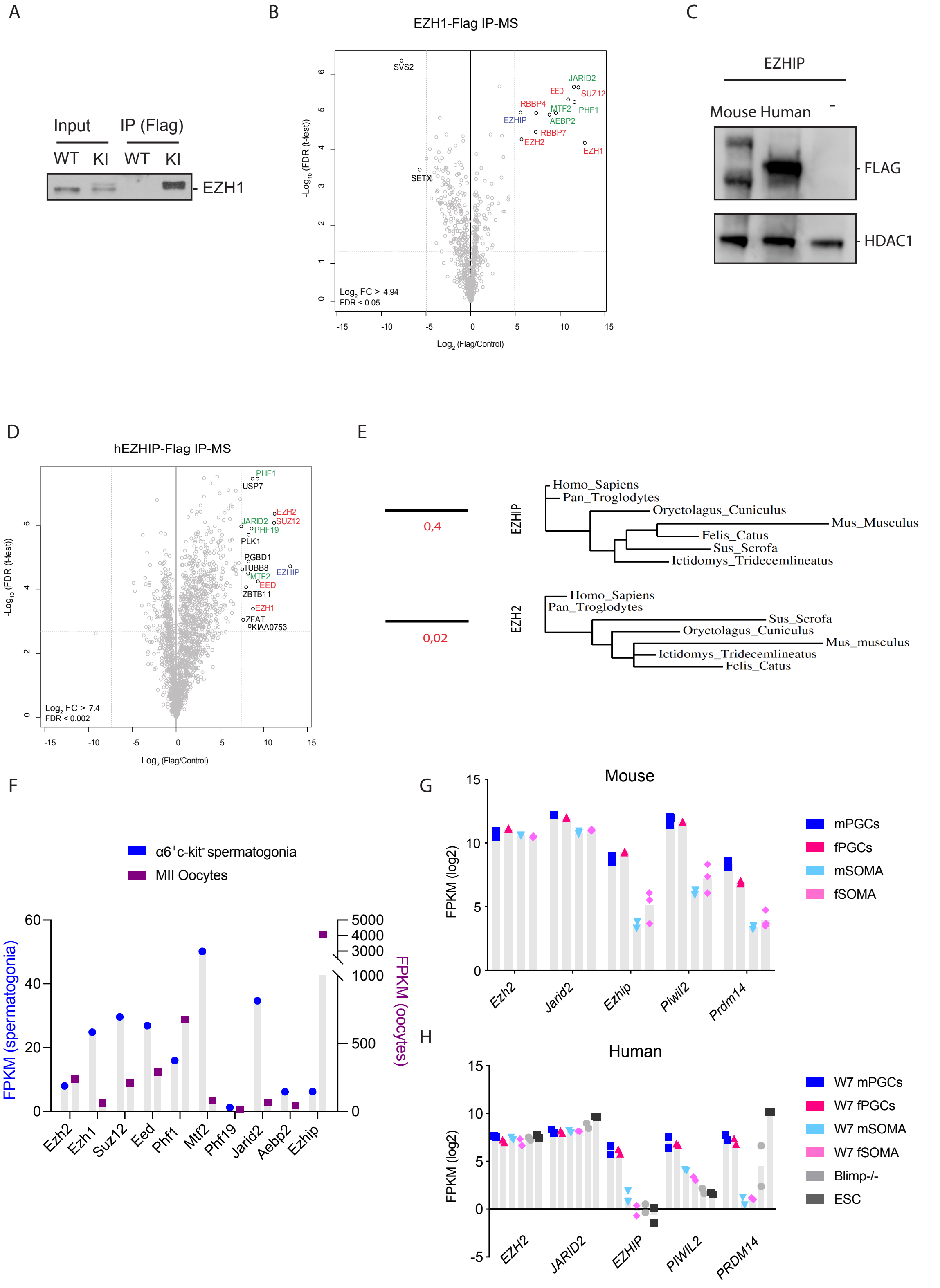

A
Input
IP (Flag)
WT
KI
WT
KI
EZH1
B
EZH1-Flag IP-MS
SVS2
JARID2
EED
SUZ12
RBBP4
MTF2
PHF1
EZHIP
AEBP2
RBBP7
EZH2
EZH1
SETX
Log2 FC > 4.94
FDR < 0.05
-Log10 (FDR (t-test))
Log2 (Flag/Control)
C
EZHIP
Mouse Human -
FLAG
HDAC1
D
hEZHIP-Flag IP-MS
PHF1
USP7
EZH2
JARID2
SUZ12
PHF19
PLK1
PGBD1
TUBB8
EZHIP
MTF2
EED
ZBTB11
EZH1
ZFAT
KIAA0753
Log2 FC > 7.4
FDR < 0.002
-Log10 (FDR (t-test))
Log2 (Flag/Control)
E
0,4
EZHIP
Homo_Sapiens
Pan_Troglodytes
Oryctolagus_Cuniculus
Mus_Musculus
Felis_Catus
Sus_Scrofa
Ictidomys_Tridecemlineatus
0,02
EZH2
Homo_Sapiens
Pan_Troglodytes
Sus_Scrofa
Oryctolagus_Cuniculus
Mus_musculus
Ictidomys_Tridecemlineatus
Felis_Catus
F
α6+c-kit- spermatogonia
MII Oocytes
FPKM (spermatogonia)
FPKM (oocytes)
Ezh2
Ezh1
Suz12
Eed
Phf1
Mtf2
Phf19
Jarid2
Aebp2
Ezhip
G
Mouse
FPKM (log2)
Ezh2
Jarid2
Ezhip
Piwil2
Prdm14
mPGCs
fPGCs
mSOMA
fSOMA
H
Human
FPKM (log2)
EZH2
JARID2
EZHIP
PIWIL2
PRDM14
W7 mPGCs
W7 fPGCs
W7 mSOMA
W7 fSOMA
Blimp-/-
ESC

**Supplementary Figure 2**

(A) Human Protein Atlas cell lines transcripts analysis (TPM: transcript per million). (B) Western blot analysis of H3K27me3 and H3 (loading control) on 293T nuclear extract and U2OS nuclear extracts WT, *EED* -/- and *EZHIP* -/- (top panel). Bottom panel, same as above for U2OS extracts but loaded with more proteins to detect H3K27me3 in WT condition, representative result. (C) RT-qPCR to detect the overexpression of *EZHIP* mutants in U2OS (control for Fig 2C, mean, n=2). (D) Different *EZHIP* C-ter truncations were expressed in HEK-293 cells (top panel). Co-IP (Flag-IP) was analyzed by western blot with the specific antibodies indicated on the right (bottom panel) (representative result, n=2). (E) GO terms the most enriched according to lowest q-value for the 287 genes found downregulated upon deletion of *EZHIP* as shown in Fig 2D. Only GO terms with a qval≤0,05 are represented.

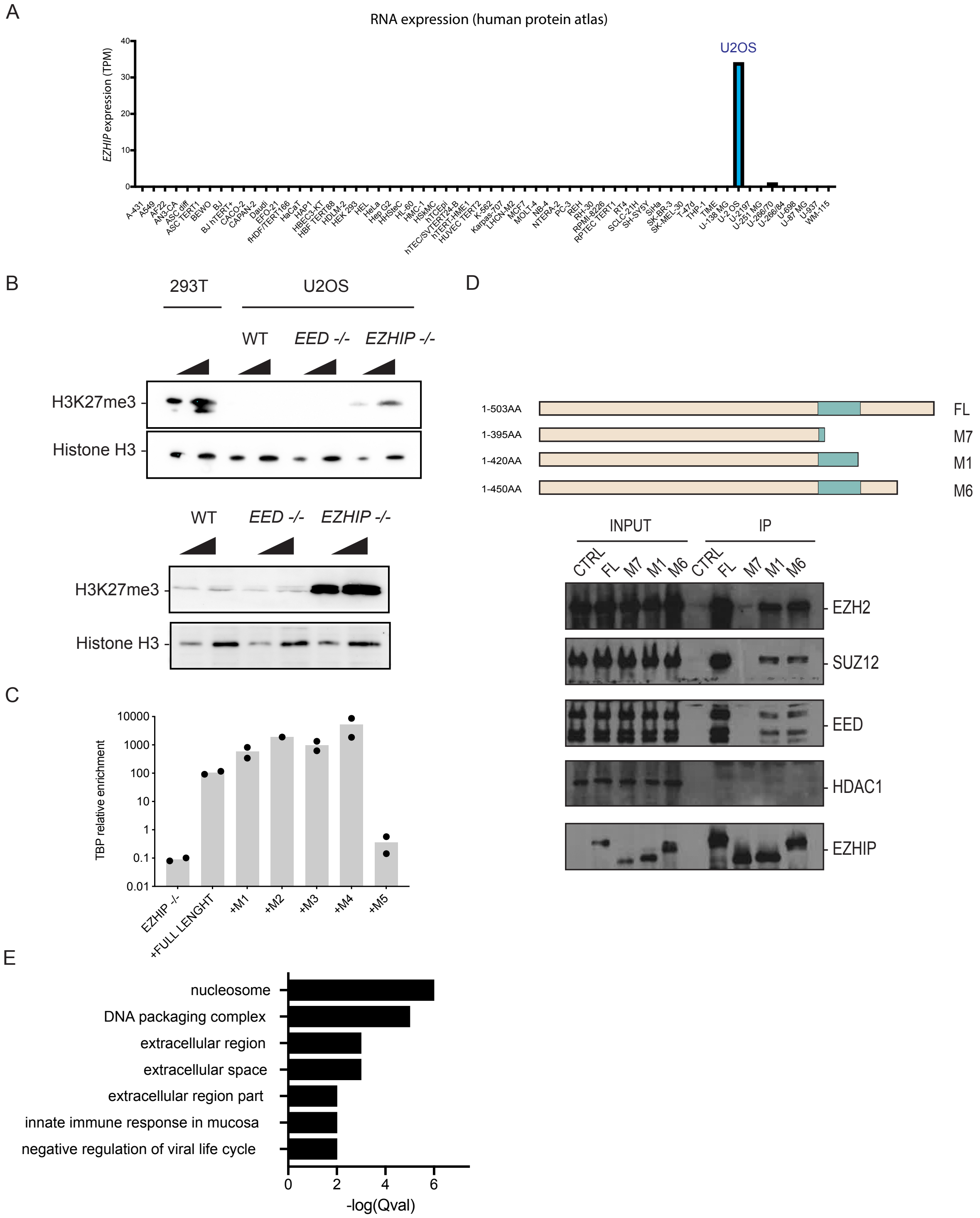
A
RNA expression (human protein atlas)
EZHIP expression (TPM)
U2OS
B
293T
U2OS
WT
EED -/-
EZHIP -/-
H3K27me3
Histone H3
WT
EED -/-
EZHIP -/-
H3K27me3
Histone H3
C
TBP relative enrichment
10000
1000
100
10
1
0.1
0.01
EZHIP -/-
+FULL LENGHT
+M1
+M2
+M3
+M4
+M5
D
1-503AA
1-395AA
1-420AA
1-450AA
FL
M7
M1
M6
INPUT
IP
CTRL
FL
M7
M1
M6
CTRL
FL
M7
M1
M6
EZH2
SUZ12
EED
HDAC1
EZHIP
E
nucleosome
DNA packaging complex
extracellular region
extracellular space
extracellular region part
innate immune response in mucosa
negative regulation of viral life cycle
0
2
4
6
-log(Qval)

**Supplementary Figure 3**

(A) Correlation heatmap for H3K27me3 ChIP-seq in U2OS WT, *EED* -/- and *EZHIP* -/- (B) Venn diagram representing the peaks detected for H3K27me3 ChIP-seq in U2OS WT *versus* U2OS *EZHIP* -/-. (C) Correlation matrix for H3K27ac, H3K27me2 and H2Aub "Cut and Run" results in U2OS WT and U2OS *EZHIP* -/-.

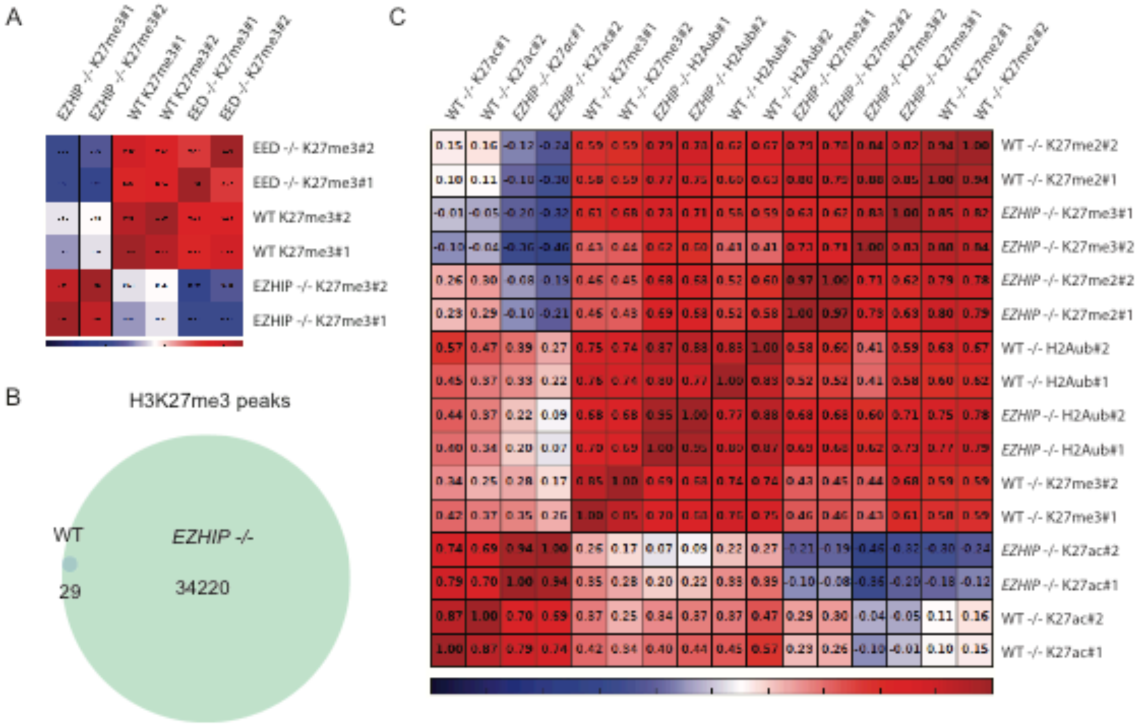

A
EZHIP -/- K27me3#1
EZHIP -/- K27me3#2
WT K27me3#1
WT K27me3#2
EED -/- K27me3#1
EED -/- K27me3#2
EED -/- K27me3#2
EED -/- K27me3#1
WT K27me3#2
WT K27me3#1
EZHIP -/- K27me3#2
EZHIP -/- K27me3#1
B
H3K27me3 peaks
WT
29
EZHIP -/-
34220
C
WT -/- K27ac#1
WT -/- K27ac#2
EZHIP -/- K27ac#1
EZHIP -/- K27ac#2
WT -/- K27me3#1
WT -/- K27me3#2
EZHIP -/- H2Aub#1
EZHIP -/- H2Aub#2
WT -/- H2Aub#1
WT -/- H2Aub#2
EZHIP -/- K27me2#1
EZHIP -/- K27me2#2
EZHIP -/- K27me3#2
EZHIP -/- K27me3#1
WT -/- K27me2#1
WT -/- K27me2#2
WT -/- K27me2#2
WT -/- K27me2#1
EZHIP -/- K27me3#1
EZHIP -/- K27me3#2
EZHIP -/- K27me2#2
EZHIP -/- K27me2#1
WT -/- H2Aub#2
WT -/- H2Aub#1
EZHIP -/- H2Aub#2
EZHIP -/- H2Aub#1
WT -/- K27me3#2
WT -/- K27me3#1
EZHIP -/- K27ac#2
EZHIP -/- K27ac#1
WT -/- K27ac#2
WT -/- K27ac#1

**Supplementary Figure 4**

(A) Heatmap representing SUZ12 enrichment in U2OS WT, *EED* -/- and *EZHIP* -/- at peaks that gain H3K27me3 enrichment in the absence of EZHIP, merged of duplicates. (B) Immunofluorescence staining for EZHIP in U2OS WT *versus* U2OS *EZHIP* -/-, nucleus stained with DAPI, representative result. (C) Left: Scheme for hEZHIP purification from Sf-9 insect cells. Right: Coomassie staining of purified protein, representative result. (D) HKMT assay performed with rPRC2-EZH2 on native nucleosomes purified from HeLa cells in presence of increasing amount of EZHIP, n≥2. (E) Flag-IPs on extracts from U2OS (WT, WT + Flag-EZH2, *EZHIP* -/- + Flag-EZH2) analyzed by western blot and probed with an antibody recognizing EZH2, representative result. (F) Volcano plot representation of normalized mass spectrometry data after Flag-IP in U2OS+ Flag-EZH2 [3], U2OS *EZHIP* -/- Flag-EZH2 (Right), same color code as in Fig.1 and S1, n=3.

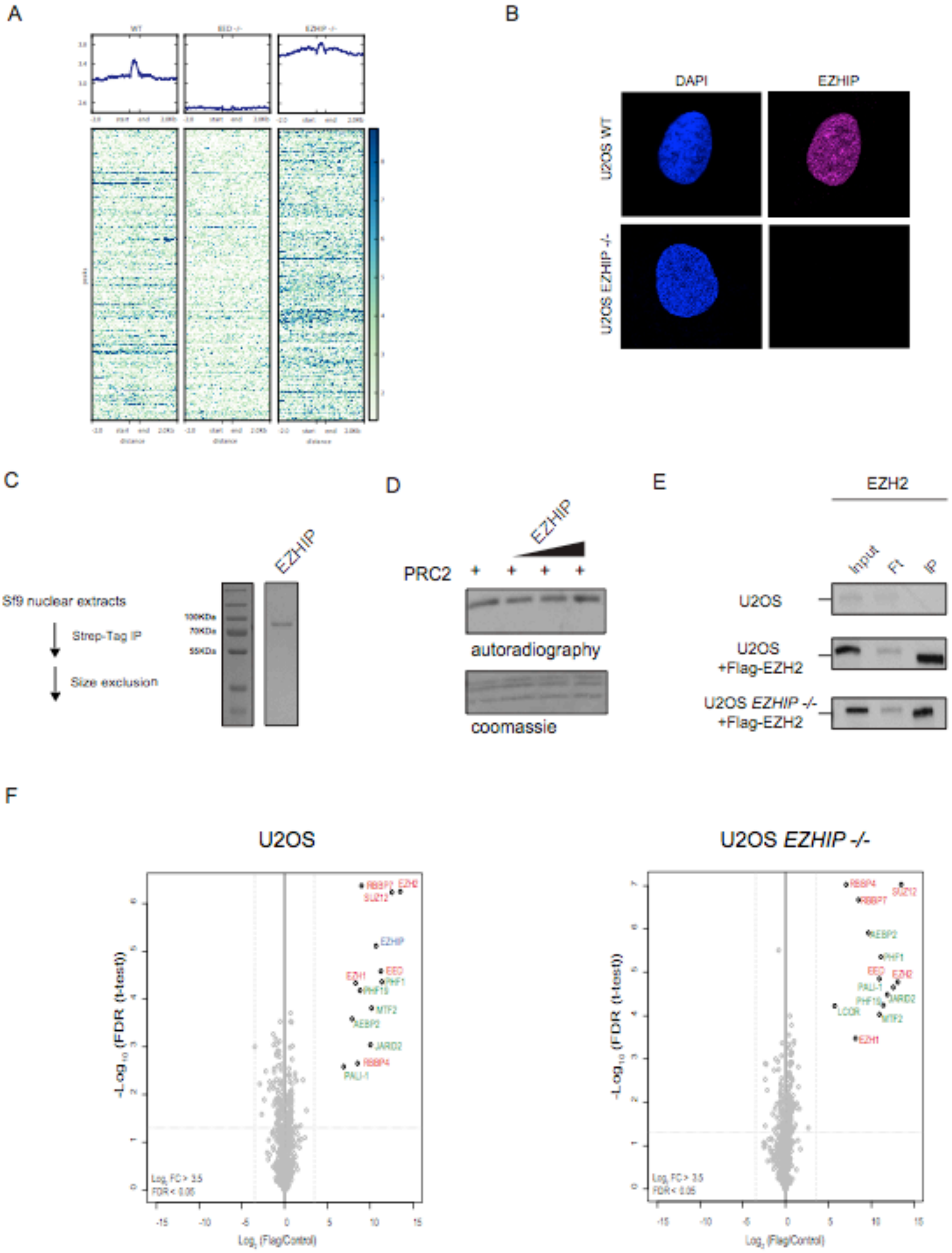
A
B
DAPI
EZHIP
U2OS WT
U2OS EZHIP -/-
C
EZHIP
Sf9 nuclear extracts
Strep-Tag IP
Size exclusion
100KDa
70KDa
55KDa
D
EZHIP
PRC2
autoradiography
coomassie
E
EZH2
Input
Ft
IP
U2OS
U2OS
+Flag-EZH2
U2OS EZHIP -/-
+Flag-EZH2
F
U2OS
U2OS EZHIP -/-
-Log10 (FDR (t-test))
Log2 (Flag/Control)
RBBP7
EZH2
SUZ12
EZHIP
EED
EZH1
PHF1
PHF19
MTF2
AEBP2
JARID2
RBBP4
PALI-1
LCOR

**Supplementary Figure 5**

(A) Schematic representation of *Ezhip* locus, scissors indicate the deletion generated by genome editing in mice. At the bottom is a representative sequencing of this locus in founders. (B) *Nudt10*, *Ezhip*, *Nudt11* and *Ezh2* mRNA relative abundancy normalized to *Gapdh* in whole testis from adult male mice WT and *Ezhip* -/-, mean, n= 3. (C) WB analysis with mouse anti-EZHIP on WT and *Ezhip* -/- testis nuclear extracts. Arrows indicate specific signal, representative result. (D) *Nudt10*, *Ezhip* and *Nudt11* mRNA normalized to *Gapdh* mRNA in whole ovaries from adult female mice WT and *Ezhip* -/-, mean, n= 3. (E) RT-qPCR analysis of *Ezhip* and *Ezh2* mRNA expression during spermatogenesis. *Ezhip* and *Ezh2* mRNA are normalized to *Tbp*. The different spermatogenic populations (undifferentiated spermatogonia kit-, differentiating spermatogonia kit+, 4N, 2N, N) have been sorted by FACS, mean, n= 2. (F) Analysis by flow cytometry of testicular cell suspensions from WT and *Ezhip* -/Y mice: spermatocyte I (4N), spermatocyte II (2N), spermatids (N), and differentiating ($\alpha$-6$^{+}$ kit$^{+}$) and undifferentiated spermatogonia ($\alpha$-6$^{+}$ kit$^{-}$). $\alpha$-6 ($\alpha$-6 integrin), kit (c-kit receptor), PI (propidium iodide), $\beta$-2m ($\beta$-2 microglubulin), and SP (Side Population) markers are indicated in graphs, representative result. (G) Mice testis absolute weight (mg), mean ±SEM, n=12. (H) Western blot analysis of H3K27me3 and H3 levels on whole testis extracts of WT; *Dnmt3l* -/- and *Ezhip* -/-; *Dnmt3 l*-/- mice. Representative result.

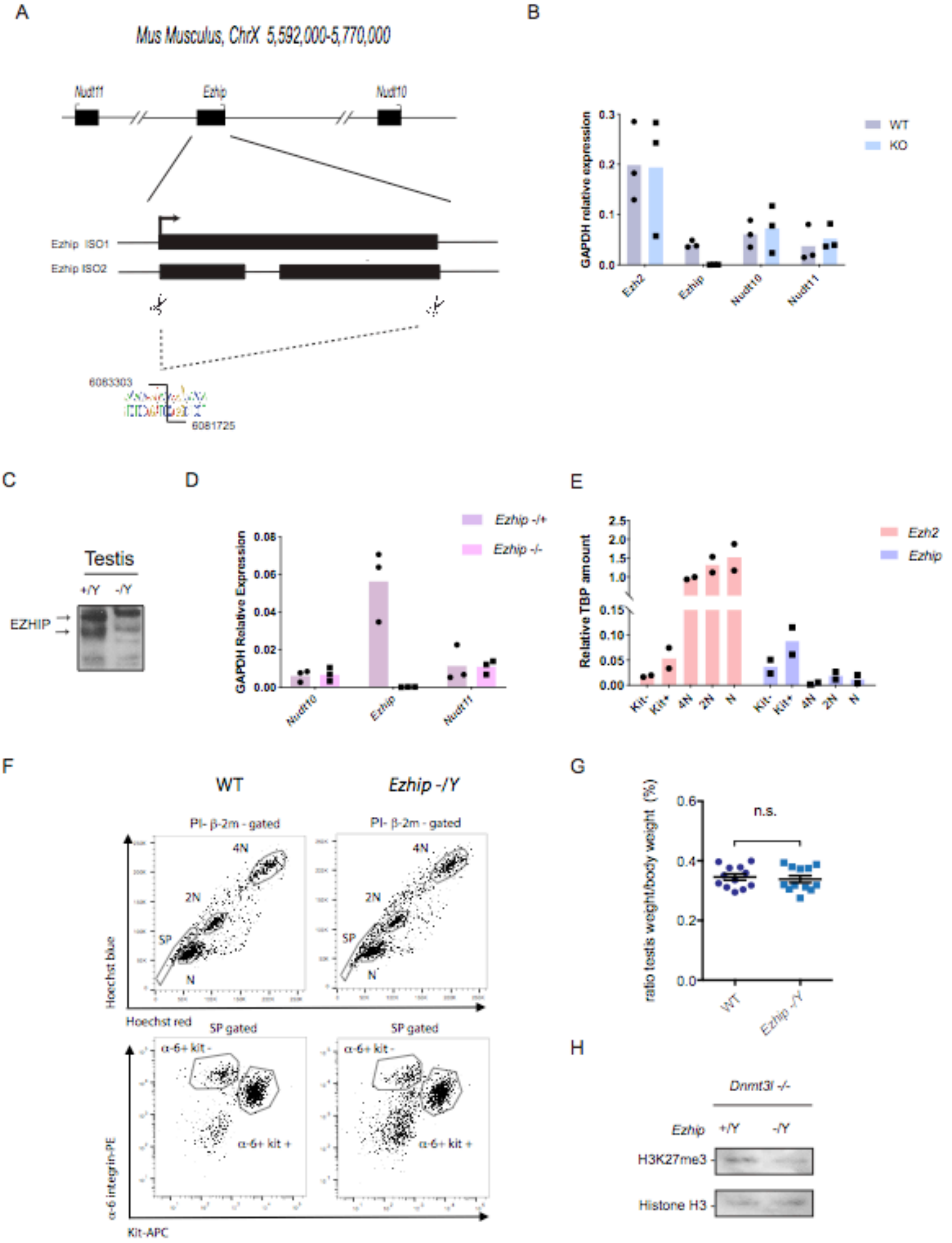

A
Mus Musculus, ChrX 5,592,000-5,770,000
Nudt11
Ezhip
Nudt10
Ezhip ISO1
Ezhip ISO2
6083303
6081725
B
GAPDH relative expression
WT
KO
Ezh2
Ezhip
Nudt10
Nudt11
C
Testis
+/Y
-/Y
EZHIP
D
GAPDH Relative Expression
Ezhip -/+
Ezhip -/-
Nudt10
Ezhip
Nudt11
E
Relative TBP amount
Ezh2
Ezhip
Kit-
Kit+
4N
2N
N
F
WT
Ezhip -/Y
PI- β-2m - gated
4N
2N
SP
N
Hoechst blue
Hoechst red
SP gated
α-6+ kit -
α-6+ kit +
α-6 integrin-PE
Kit-APC
G
ratio testis weight/body weight (%)
n.s.
WT
Ezhip -/Y
H
Dnmt3l -/-
Ezhip
+/Y
-/Y
H3K27me3
Histone H3

**Supplementary Figure 6**

(A) Non-surrounded Nuclei (NSN) WT and *Ezhip* -/- oocytes were fixed and stained for H3K27me3 (green in merged). DNA was stained with DAPI (blue in merged). IF quantification is indicated on the right, *n* number of oocytes analyzed, H3K27me3 intensities are normalized to DAPI. (B) Same as in (A) but Surrounded Nuclei (SN) oocytes were probed for H3K4me3.

A

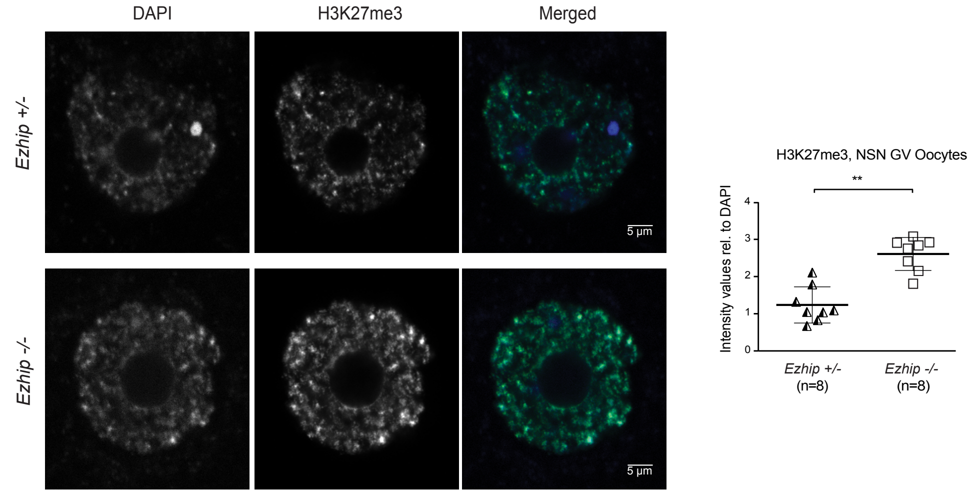
DAPI
H3K27me3
Merged
Ezhip +/-
Ezhip -/-
5 µm
5 µm
H3K27me3, NSN GV Oocytes
**
Intensity values rel. to DAPI
4
3
2
1
0
Ezhip +/-
(n=8)
Ezhip -/-
(n=8)

B

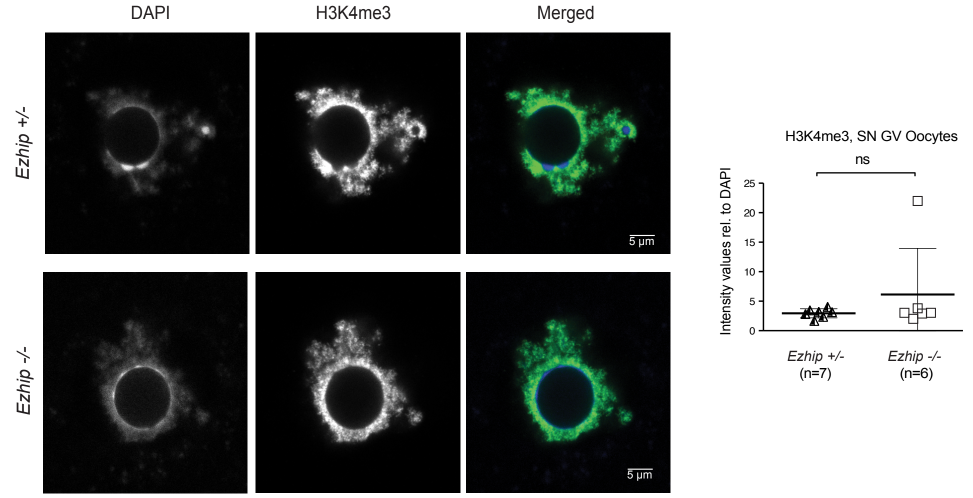
DAPI
H3K4me3
Merged
Ezhip +/-
Ezhip -/-
5 µm
5 µm
H3K4me3, SN GV Oocytes
ns
Intensity values rel. to DAPI
25
20
15
10
5
0
Ezhip +/-
(n=7)
Ezhip -/-
(n=6)

**Supplementary Figure 7**

Images showing representative reproductive track of 6-weeks-old WT and *Ezhip* -/- females.

WT *EZHIP -/-*

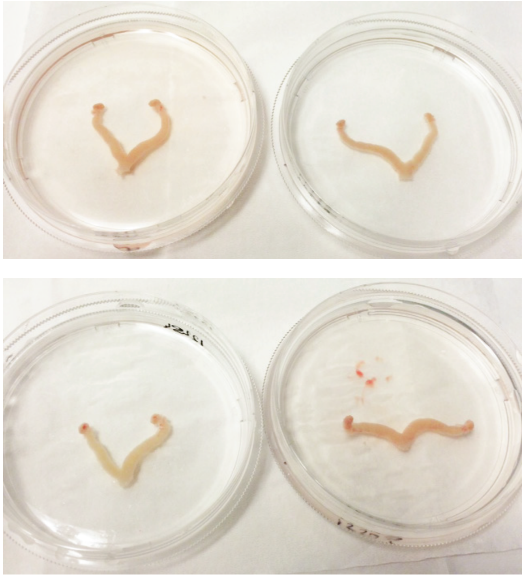

### Supplementary Table 1

Full analysis of computer-assisted spermatozoa images.
In red are the numbers represented in Fig 5E.

| General Characteristics | KOY | KOY | WTY | WTY |
|---|---|---|---|---|
| Path velocity | 140,8 | 147,5 | 135,5 | 132,5 |
| Prog. Velocity | 109 | 110,6 | 106,7 | 97,8 |
| Track speed | 251,3 | 280,6 | 243,6 | 255 |
| Lateral amplitude | 10,3 | 11,4 | 10 | 10,9 |
| Beat frequency | 28,2 | 28,8 | 28,4 | 30,6 |
| Stainghtness | 77 | 74 | 77 | 74 |
| Linearity | 47 | 41 | 47 | 40 |
| Elongation | 53 | 54 | 53 | 51 |
| Area | 47,1 | 48,7 | 47,3 | 54 |
| **Sperm properties** | | | | |
| Total sperm cells counted | 428 | 868 | 595 | 1311 |
| Motile | 259 | 631 | 426 | 1140 |
| Progressive | 143 | 311 | 249 | 525 |
| Expressed as Percentage of the total % | 100 | 100 | 100 | 100 |
| | 61 | 73 | 72 | 87 |
| | 43 | 36 | 42 | 42 |
| **Sperm speed** | | | | |
| Rapid | 243 | 599 | 415 | 1096 |
| Medium | 16 | 32 | 11 | 44 |
| Slow | 3 | 4 | 4 | 8 |
| Static | 166 | 233 | 165 | 163 |
| Expressed as Percentage of the total % | 57 | 69 | 70 | 84 |
| | 4 | 4 | 2 | 3 |
| | 1 | 0 | 1 | 1 |
| | 39 | 27 | 28 | 12 |

### Supplementary Table 2

Primers sequences.

| Name | Application | Sequence |
|---|---|---|
| mGapdh FW | RT-qPCR | AACAGCAACTCCCACTCTTC |
| mGapdh REV | RT-qPCR | TGGTCCAGGGTTTCTTACTC |
| mEzh2 FW | RT-qPCR | AATACATGTGCAGCTTTCTGTTC |
| mEzh2 REV | RT-qPCR | ACGAATTTTGTTGCCCTTTC |
| mEzhip FW | RT-qPCR | TTCCGGAGTTGTACCTTTCG |
| mEzhip REV | RT-qPCR | ACGTAAATTCCAGCCTGTGC |
| mNudt10 FW | RT-qPCR | AGAGAGCGAGCCCTAGTGAATGGA |
| mNudt10 REV | RT-qPCR | GAGCTCACCTGTGCTTCACAATTCC |
| mNudt11 FW | RT-qPCR | ACCGAGGCATGCTCAAGATCACA |
| mNudt11 REV | RT-qPCR | TGAGCGGTCTCCTTGGCAACCTTA |
| hEZHIP N-ter FW | RT-qPCR | ACCTCCGCCGCCATTTTCATCA |
| hEZHIP N-ter REV | RT-qPCR | TCGGGCACCACACACCCAAAAA |
| hEZHIP stretch FW | RT-qPCR | GCCTGTTTGGCATGCAGTCCGTAT |
| hEZHIP stretch REV | RT-qPCR | ACTGCTGAGGGATGGGAAGGAAGA |
